# Personalised Regional Modelling Predicts Tau Progression in the Human Brain

**DOI:** 10.1101/2023.09.28.559911

**Authors:** Pavanjit Chaggar, Jacob Vogel, Alexa Pichet Binette, Travis B. Thompson, Olof Strandberg, Niklas Mattsson-Carlgren, Linda Karlsson, Erik Stomrud, Saad Jbabdi, Stefano Magon, Gregory Klein, the Alzheimer’s Disease Neuroimaging Initiative, Oskar Hansson, Alain Goriely

**Affiliations:** Mathematical Institute, University of Oxford; Clinical Memory Research Unit, Department of Clinical Sciences Malmo, Lund University; Department of Mathematics and Statistics, Texas Tech University, USA; Wellcome Trust Centre for Integrative Neuroscience, University of Oxford; F. Hoffmann-La Roche Ltd, Basel, Switzerland; Memory Clinic, Skane University Hospital, Lund University; Department of Neurology, Skåne University Hospital, Lund University; Wallenberg Center for Molecular Medicine, Lund University

## Abstract

Aggregation of the hyperphosphorylated tau protein is a central driver of Alzheimer’s disease, and its accumulation exhibits a rich spatio-temporal pattern that unfolds during the course of the disease, sequentially progressing through the brain across axonal connections. It is unclear how this spatio-temporal process is orchestrated – namely, to what extent the spread of pathologic tau is governed by transport between brain regions, local production or both. To address this, we develop a mechanistic model from tau PET data to describe tau dynamics along the Alzheimer’s disease timeline. Our analysis reveals longitudinal changes in production and transport dynamics on two independent cohorts, with subjects in early stage of the disease exhibiting transport-dominated spread, consistent with an initial spread of pathologic tau seeds, and subjects in late stage disease (Braak stage 3/4 onwards) characterised primarily by local production of tau. Furthermore, we demonstrate that the model can accurately predict subject-specific longitudinal tau accumulation at a regional level, potentially providing a new clinical tool to monitor and classify patient disease progression.

**Teaser:** A mechanistic model reveals tau protein dynamics in Alzheimer’s, showing stage-specific shifts in transport and local production.

## 1 Introduction

Alzheimer’s disease (AD) is a devastating neurological condition resulting in progressive brain atrophy and cognitive decline. Toxic forms of two proteins, amyloid-*β* (A*β*) and tau protein (*τ*P) are believed to act in concert to drive AD progression [1, 2]. The pathological roles of these proteins in the human brain during AD has been investigated using positron emission tomography (PET), with radiotracers such as [^18^F]florbetapir and [^18^F]flortaucipir allowing for in-vivo quantification of A*β* and *τ*P, respectively [3]. While A*β* tends to be more diffusely present throughout the cerebral cortex [4, 5, 6, 7], *τ* P exhibits richer spatiotemporal dynamics, characterised by Braak staging [8]. Braak staging describes the trajectory of toxic *τ* P, starting from the entorhinal cortex and progressing sequentially into the limbic regions, basal temporal lobes, broad association cortex and finally into primary sensory cortex. This staging pattern has been validated using *τ*P-PET imaging [9, 10, 11] and has been shown to be highly correlated with atrophy and cognitive decline [12, 13], however, the mechanism for how *τ*P staging is orchestrated remain unclear.

Growing evidence suggests that the progression of AD depends on two key factors: 1) the transport of toxic proteins throughout the brain; 2) the local production of toxic proteins. However, the extent to which these factors contribute to AD progression and whether their contributions change over time has yet to be determined. There is now substantial evidence that *τ*P propagation follows a prion-like mechanism, progressively forming toxic oligomeric seeds and neurofibrillary tangles through an autocatalyic production process [14, 15]. The prion-like nature of *τ*P has been demonstrated with transgenic animal models in which cortical injections of *τ*P seeds induces the formation of *τ*P aggregates that grow in concentration over time at the site of injection and surrounding areas [16, 17]. In 2012, studies from Liu et al. and de Calignon et al. showed that transgenic mice overexpressing pathological human *τ*P in the entorhinal cortex exhibit accumulation of *τ*P aggregates and that *τ*P invades axonally connected regions through trans-synaptic transport to form seeds in otherwise healthy regions [18, 19]. The prion-like aggregation and axon-based transport of *τ*P has also been suggested in human post-mortem studies [20] and by in-vivo studies using structural connectome-based models of *τ*P PET capable of reproducing observed *τ*P aggregation and spreading [21, 22, 23, 24, 25, 26]. In a recent investigation, Meisl et al. analysed multimodal *τ*P data from Braak stage 3 onward and showed that *τ*P production, not transport, is the main contributor of *τ*P progression [27]. However the study does not account for the spatial progression across individual brain regions or estimate dynamics across the full AD progression timeline. It remains unanswered whether there are changes in the rates of *τ*P production and transport over time and whether the balance of these two processes change along the disease timeline. To address these outstanding questions, we develop a whole-brain model capable of accurately describing longitudinal *τ*P PET data and conduct a multi-cohort study to analyse *τ*P dynamics across the full disease progression timeline.

To answer questions about temporal changes in AD *τ*P dynamics in the human brain requires an accurate and reliable model of longitudinal *τ*P observations. There have been numerous efforts over the past decade to use mathematical models to better understand the spatiotemporal properties of AD pathology, ranging from linear diffusion models of *τ*P [21, 22] to infinite dimensional spatiotemporal models of toxic protein aggregation [28]. Each of these models make different assumptions about the physical mechanisms of *τ*P spread, however, there has not been a unifying effort to rigorously compare commonly used models to identify which are best able to accurately describe longitudinal *τ*P PET observations. In addition, the models currently described in the literature fail to account for regional variations in *τ*P dynamics, which has been shown to influence the progression of *τ*P [29, 30, 31] and are incapable of predicting longitudinal changes at a regional level. Here, we present a novel model that provides a qualitative account of regional vulnerability and its effect on *τ*P progression. Using a previously developed Bayesian pipeline for longitudinal *τ*P modelling [32, 26], we perform hypothesis-driven model selection on a family of common models from the AD modelling literature, including a new model accounting for regional dynamics. We show that models relying only on network diffusion or homogeneous *τ*P production dynamics are not sufficient to model regional longitudinal data, whereas models accounting for regional variations in *τ*P dynamics are able to accurately model longitudinal *τ*P observations at a regional level.

We combine state-of-the-art modelling and inference methods with longitudinal *τ*P PET data from two independent cohorts to address the outstanding question of how the transport and production of *τ*P drive AD progression. To determine whether there are changes in the rates of *τ*P transport and production during AD progression, we apply our model to three groups A*β*^*−*^, A*β*^+^/*τ*P^*−*^ and A*β*^+^/*τ*P^+^ subjects, representing different stages of the AD timeline [33]. We show that the transport of *τ*P is faster in early stage disease (A*β*^+^/*τ*P^*−*^), and that there are primary and secondary increases in production dynamics of *τ*P along the disease timeline. We validate these results on an independent dataset using a different *τ*P tracer, BioFINDER-2, on which the same results are obtained, further showing that the model and results are robust and generalisable across datasets and choice of *τ*P tracer. Finally, we validate our model by showing that it can *predict* regional rates of *τ*P accumulation over time for individual patients. The combination of these methods provide a novel pipeline for analysing and understanding longitudinal *τ*P data, allowing for us to compare changes in disease dynamics across the AD timeline and predict subject specific, region specific changes in *τ*P over time.

## 2 Results

### 2.1 Deriving a generative model of *τ*P dynamics

We first extend previous work [34, 35, 28] to develop a mechanism-based model of *τ*P dynamics in the human brain that can be calibrated using *τ*P PET data. This model, called the *local FKPP model* (after the well known Fisher-Kolmogorov–Petrovsky–Piskunov equation) and derived in full detail in Section 4.4, is given by a set of nonlinear ordinary differential equations for the variables *s*_*i*_ = *s*_*i*_(*t*) representing *τ*P SUVR in different regions of interest on a connectome with *N* nodes:

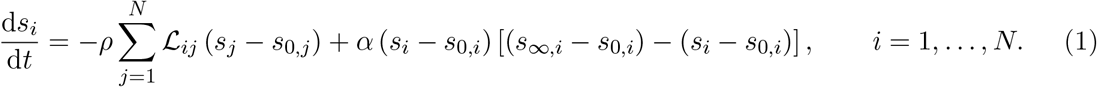

The first term represents transport of *τ*P between brain regions through a graph Laplacian ℒ with uniform rate *ρ*, and is consistent with previous work [21, 22, 24, 35]. In this study, regions of interest are given by cortical regions of the Desikan-Killiany-Tourville (DKT) atlas, and the bilateral hippocampus and amygdala (N = 72). We introduce two novel parameter vectors, regional baseline values, *s*_0,*i*_, and carrying capacities *s*_*∞,i*_, that represent a healthy state and a late-stage AD state, respectively, which add information about regional variations in production dynamics. These parameters are estimated using the Gaussian mixture modelling approach developed in [24] (see Section 4.5 for details). An example of this is shown in Fig. 1a for the right inferior temporal lobe and the carrying capacities for the right hemisphere are shown in Fig. 1b. Since these parameters are estimated from tau PET, they also encode specific features of the tracer, such as regional differences in tracer uptake, specificity to 3R/4R tau pathology, on-target binding and off-target binding, and allows us to model tau SUVR directly. The simulated transport and production dynamics are shown in Fig. 1. A consequence of the variation of carrying capacities is that the regional production rates also vary between regions, as seen in the time series of Fig. 1c, providing a qualitative account of regional vulnerability. The ability to account for these regional variations extends previous models with homogeneous dynamics across regions [35, 23] provides a picture of *τ*P progression that is more consistent with observed *τ*P staging.

**Figure 1.**
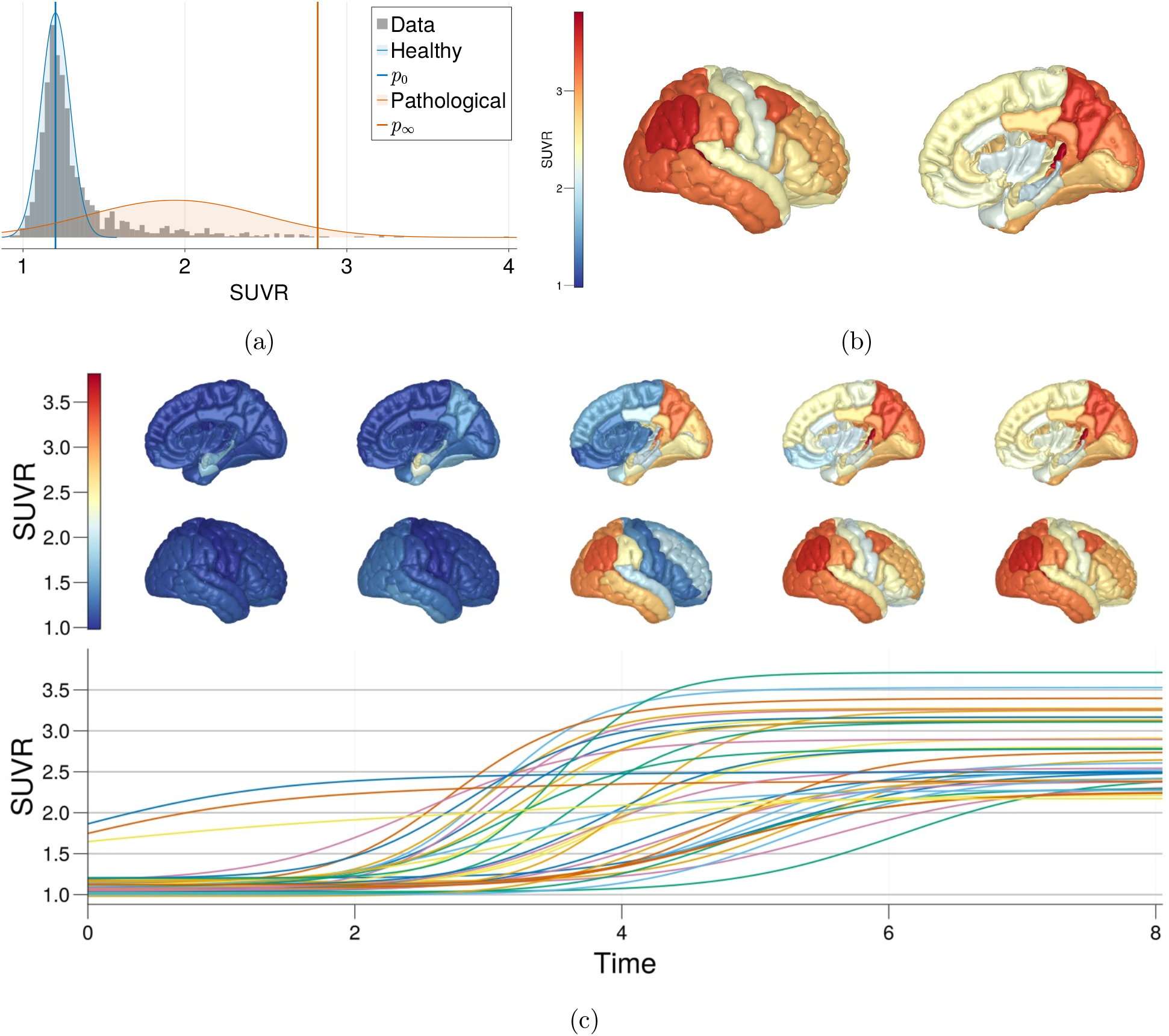
Simulated transport and production dynamics in the local FKPP model. **1a** Two component Gaussian mixture model fit to a multi-cohort *τ*P PET dataset [24] and data from ADNI for the right inferior temporal lobe. Baseline values are taken as the mean of the *τ*P^*−*^ distribution and the carrying capacity as the 99-th percentile of the *τ*P^+^ distrbiution. **1b** Right hemisphere cortical rendering of the SUVR carrying capacities. **1c** Simulation from the local FKPP model using carrying capacities derived from Gaussian mixture models. Simulations are initialised with a seed of 50% concentration in the bilateral medial temporal lobe (entorhinal cortex, hippocampus and amygdala), with *ρ* = 0.15 and *α* = 1.1. Each line represents the SUVR trajectory of one DKT atlas brain region. Values at time points *t* = {0, 2, 4, 6, 8} are projected onto a cortical rendering.

### 2.2 Regional Heterogeneity Is Necessary for Longitudinal Prediction

To determine whether the local FKPP is capable of fitting observed AD trajectories, we compare it to *τ*P PET data. For comparison, we also consider simpler models that can be obtained from the local FKPP model Eq. (1), namely *the global FKPP model*, Eq. (10), obtained by assuming that none of the parameters vary locally, *the diffusion model* Eq. (5) obtained by taking *α* = 0 in Eq. (1), and *the logistic model* obtained by neglecting transport between regions (*ρ* = 0 and given explicitly by Eq. (12)).

We use hierarchical Bayesian inference to calibrate each model to *τ*P PET data, allowing us to quantify whether the model parameters can be identified from patient data and provides bounds on uncertainty for group and individual level parameters. We use *τ*P PET data from ADNI, selecting A*β*^+^ subjects who have at least three scans and are *τ*P^+^ in the medial temporal or lateral temporal lobe (see Section 4.1 for details). We employ two metrics to compare models, the maximum likelihood value (*L*_*max*_) to measure in-sample fit and the expected log predictive density (ELPD) to measure out-of-sample predictive accuracy, both provided in Table 1. Fig. 2 shows the in-sample longitudinal fit for each of the four models. The out-of-sample fit for the local FKPP model is shown in Fig. 4 and and in Fig. S1 for the remaining models.

**Table 1:** Assessment of model fit using the maximum log-likelihood *L*_*max*_ and the expected log predictive density; higher values correspond to better models. The local FKPP model performs best in both metrics.

|  | Local FKPP | Global FKPP | Diffusion | Logistic |
| --- | --- | --- | --- | --- |
| $L_{max}$ | **10279.40** | 9357.79 | 6635.11 | 9873.54 |
| ELPD | **300.10** | 67.12 | -450.00 | 261.84 |

**Figure 2.**
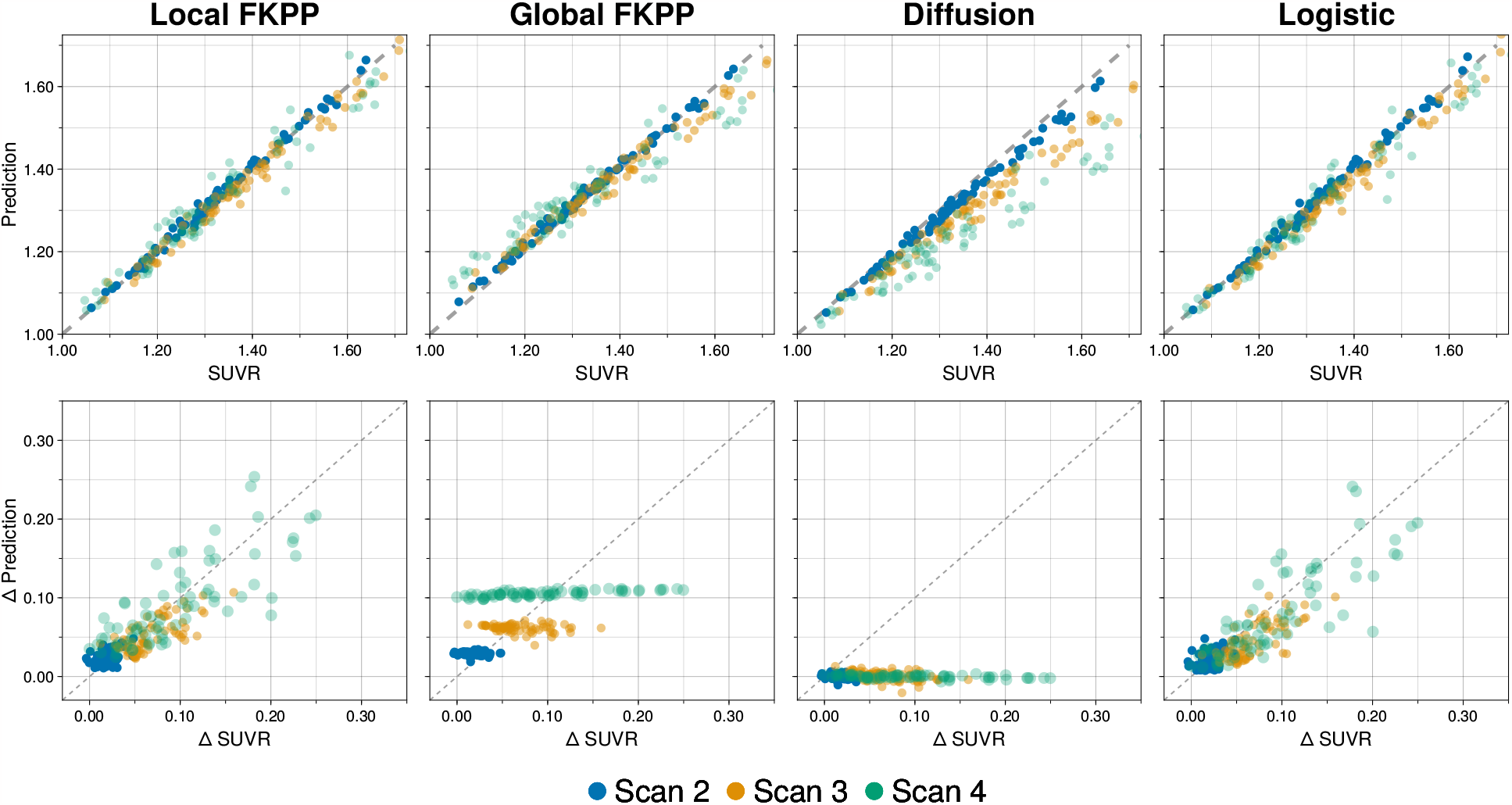
Goodness of fit for four models, local FKPP, global FKPP, diffusion and logistic. For all panels, each point represents a region in the connectome model, averaged over subjects per scan. Top row shows estimated vs observed SUVR values. Bottom row shows estimated change vs observed change in SUVR. Only the local FKPP and logistic model are able to accurately capture longitudinal changes, while the global FKPP and diffusion models are each structurally incapable of describing heterogeneous production.

The local FKPP performs best for both in-sample fit and out-of-sample predictive accuracy, closely followed by the logistic model. The results show that the global FKPP and diffusion models are ill-equipped for longitudinal modelling of *τ*P PET data, shown clearly in Fig. 2 and Fig. S1. The diffusion model underestimates SUVR evolution in time since there is no mechanism for *τ*P production or clearance and therefore total concentration is conserved. The global FKPP model is able to capture the changes in *τ*P load, but it cannot describe the regional heterogeneity in *τ*P production. The deficiencies in production dynamics of the diffusion and global FKPP models are addressed with the local FKPP model and the logistic model and the ability of these models to accurately describe longitudinal *τ*P PET underlines the importance of including regionally specific production rates. The model selection results support the role of *τ*P production playing a dominant role in driving AD pathology. The logistic model is able to describe the trajectory of *τ*P PET despite not being able to capture the transport of *τ*P through the structural connectome. This could either suggest that A*β*^+^/*τ*P^+^ subjects already have widespread invasion of *τ*P seeds or that the diffusion of *τ*P occurs over a very long timescale and the effects are not prominent in the relatively short time window over which longitudinal ADNI data is available. The error in model fit relative to the final in-sample scan is shown in Fig. S3 for the local FKPP and logistic model and show that the logistic model is prone to overestimated the SUVR (albeit very slightly), whereas the error in the local FKPP fit is more balanced across regions, possibly due to the effect of transport between regions. Overall, the data support the use of local FKPP model, evidenced by it being the most capable at describing in-sample and out-of-sample data, while also capturing the role of both tau transport and local tau production.

### 2.3 Early AD progression is driven by *τ*P transport

We next sought to determine whether there are any changes in *τ*P production and transport dynamics across the AD progression timeline. To do so, we use two cohorts of *τ*P PET data, ADNI and BioFINDER-2 (BF2), each divided into three groups, A*β*^*−*^, A*β*^+^/*τ*P^*−*^, A*β*^+^/*τ*P^+^, representing different stages of AD. Since BF2 uses a different *τ*P PET radiotracer, we rerun the Gaussian mixture modelling analysis to recover tracer specific baseline and carrying capacities. We then fit a hierarchical Bayesian model to each of the groups and examine the population level production and transport parameters.

The population parameter distributions for the A*β*^*−*^, A*β*^+^/*τ*P^*−*^, A*β*^+^/*τ*P^+^ groups are shown in Fig. 3a for ADNI and Fig. 3b for BF2 and summarised in Table 2. The posterior distributions across cohorts are qualitatively the same, with changes likely reflecting differences in cohort and tracers. The inferred parameters show an increase in the transport rate for the A*β*^+^/*τ*P^*−*^ group relative to the A*β*^+^/*τ*P^+^ and A*β*^*−*^ groups, suggesting that *τ*P more readily spreads between regions during early stages of disease, and is minimal in later stages of AD. The inferred posterior distributions for the production parameter show a progressive increase in production rate along the disease timeline, with a primary increase from A*β*^*−*^ to A*β*^+^/*τ*P^*−*^ group and a secondary increase from the A*β*^+^/*τ*P^*−*^ group to the A*β*^+^/*τ*P^+^ group. The negative production rate for the A*β*^*−*^ group indicates that the signal, on average, decreases. This could reflect changes in noise due to off-target, nonspecific binding or atrophy from non-AD related neurodegeneration. Therefore, any observed clearance dynamics likely reflect fluctuations in noise or age-related atrophy. These results that in early AD, *τ*P begins in a transport dominated phase (*ρ > α*) and later switches to a production dominated phase (*α > ρ*).

**Table 2:** Posterior summary providing the means of inferred population parameters for ADNI and BF2.

| **Group** | | $\rho_\mu$ | | $\rho_\sigma$ | | $\alpha_\mu$ | | $\alpha_\sigma$ | |
| --- | --- | --- | --- | --- | --- | --- | --- | --- | --- |
|  |  | mean | s.d. | mean | s.d. | mean | s.d. | mean | s.d. |
| ADNI | $A\beta^+/\tau P^+$ | 0.01 | 0.003 | 0.02 | 0.003 | 0.12 | 0.02 | 0.11 | 0.02 |
| | $A\beta^+/\tau P^-$ | 0.04 | 0.01 | 0.15 | 0.03 | -0.005 | 0.05 | 0.26 | 0.04 |
| | $A\beta^-$ | 0.02 | 0.01 | 0.05 | 0.01 | -0.05 | 0.03 | 0.19 | 0.02 |
| BF2 | $A\beta^+/\tau P^+$ | 0.004 | 0.001 | 0.01 | 0.001 | 0.11 | 0.01 | 0.09 | 0.01 |
| | $A\beta^+/\tau P^-$ | 0.02 | 0.01 | 0.02 | 0.01 | 0.02 | 0.05 | 0.22 | 0.04 |
| | $A\beta^-$ | 0.02 | 0.01 | 0.08 | 0.01 | -0.03 | 0.03 | 0.19 | 0.03 |

**Figure 3.**
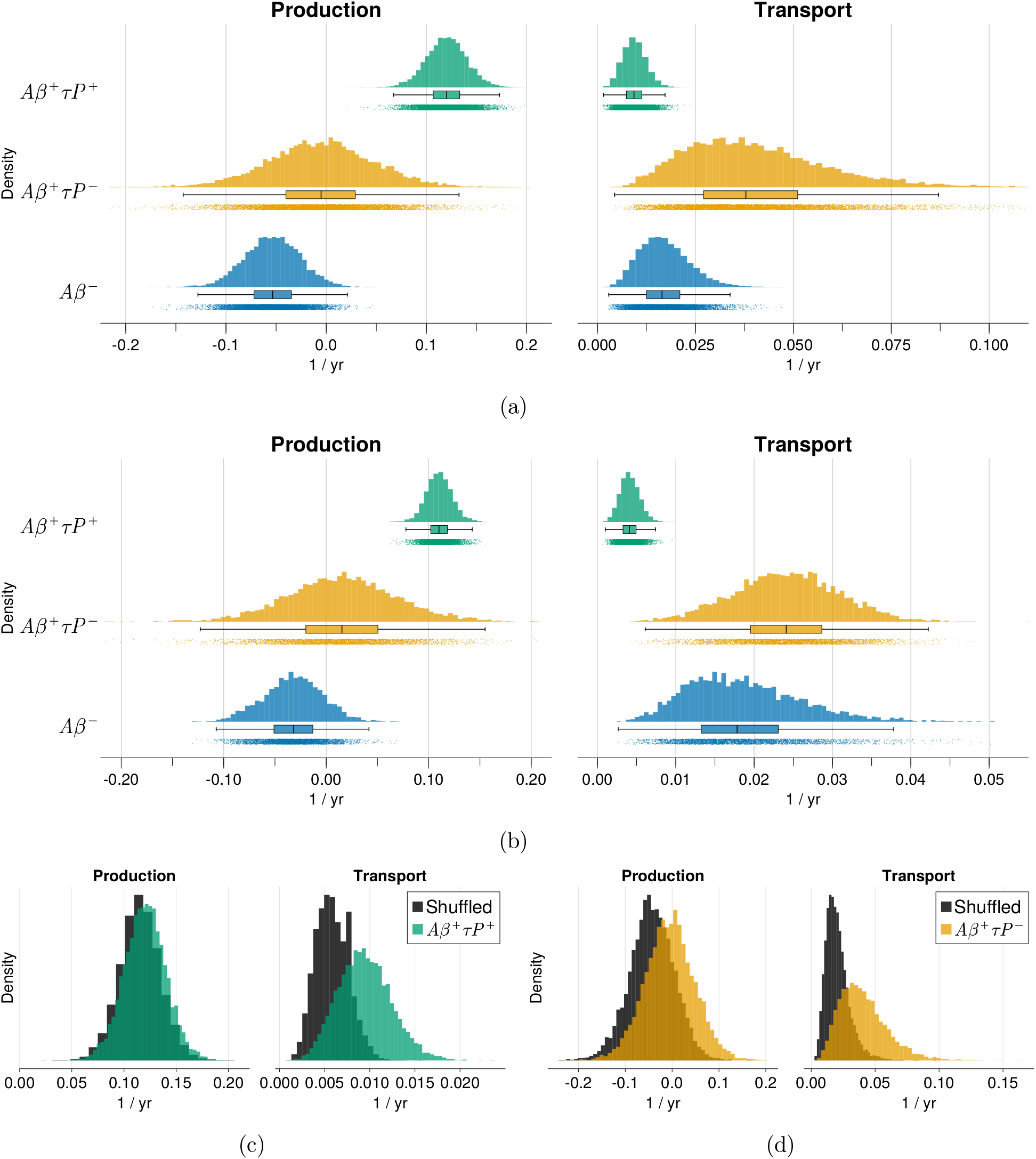
Inferred population level parameters using ADNI and BF2 *τ*P data. **3a & 3b** Population production and transport parameters across A*β*^+^/*τ*P^+^, A*β*^+^/*τ*P^*−*^ and A*β*^*−*^ groups for ADNI (3a) and BF2 (3b) *τ*P PET data. **3c & 3d** Inferred population production and transport parameters from spatially shuffled data (shown in grey) for A*β*^+^/*τ*P^+^ (3c), A*β*^+^/*τ*P^*−*^ (3d) ADNI groups.

**Figure 4.**
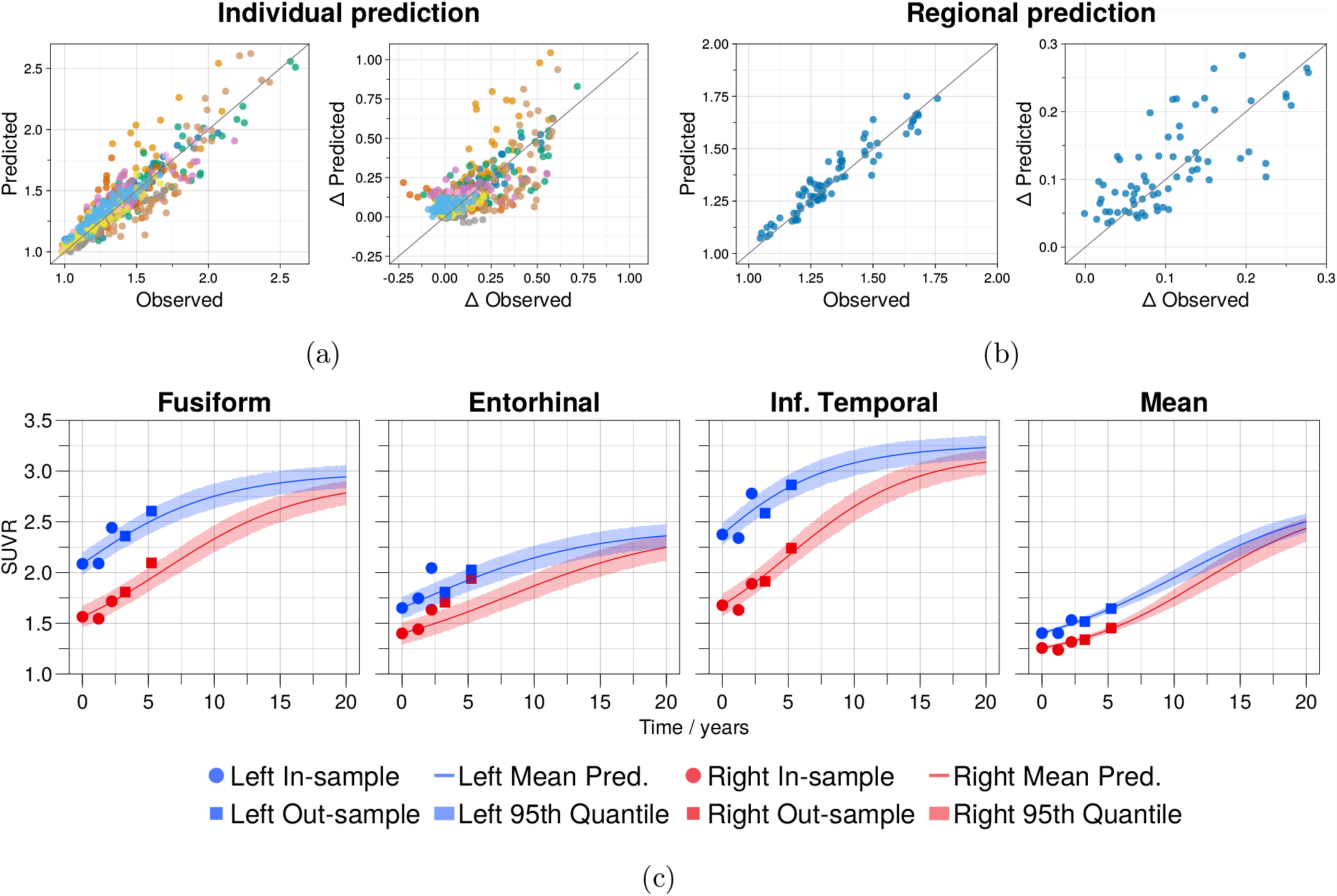
Out-of-sample fit and regional posterior predictive plots. **4a** Predicted vs observed out-of-sample SUVR for all ten subjects used for out of sample prediction. Each color represents a different patient and each point of a given color represents a region. **4b** Predicted vs observed SUVR per region, averaged over subject. Each point represents a region in the DKT atlas. **4c** Predicted regional changes in SUVR for a subject with five scans. The model is trained on the first three observed scans (circles), with the last two scans (squares) left out. We show three regions of the DKT atlas, the right fusiform, entorhinal cortex and inferior temporal lobe and the mean SUVR. 2000 simulations were run using values from the posterior distributions. Predictions for the remaining subjects who have greater than three scans are provided in Fig. S2.

To confirm that the parameter distributions reflect meaningful dynamics present in the data and are not as a result of statistical patterns, we re-run the analysis on the A*β*^+^/*τ*P^*−*^ and A*β*^+^/*τ*P^+^ groups using spatially shuffled data. The results of these for the production and transport parameters are shown in Figs. 3c and 3d. Note that we only spatially shuffle the data and therefore expect minimal changes to the estimated production parameters. We note a marked difference in estimated transport dynamics in both the A*β*^+^/*τ*P^*−*^ and A*β*^+^/*τ*P^+^ groups between the true and shuffled data, confirming that dynamics present in the data are not a consequence of statistical patterns in the data but represent *τ*P dynamics as measured through PET. The transport dominated phase of the early AD subjects supports evidence showing *τ*P seeds are present throughout the cortex before symptom onset [20, 36] and, together with the small role of transport in the A*β*^+^/*τ*P^+^ groups, helps explain the strong performance of the logistic model in Section 2.2. Overall the results reveal temporal changes in the dynamics of *τ*P progression, with an initial transport dominated phase, perhaps in which seeds are deposited around the cortex, followed by a production dominated phase indicative of secondary tauopathy, likely due to spatial colocalisation with A*β* catalysing *τ*P production.

### 2.4 Local model predicts regional *τ*P progression

A major utility of mathematical modelling of AD lies in the application to pharmacological and clinical research, particularly through predicting patient disease progression. Here, we show that the local FKPP model can be used to accurately *predict* the trajectory of *τ*P PET. Taking A*β*^+^/*τ*P^+^ subjects from ADNI who have at least four scans (*N* = 10), we use the first three scans for calibrating the model and the remaining scans for prediction. The results of this for the local FKPP model are displayed in Fig. 4 and show that the model is capable of predicting longitudinal regional *τ*P PET data. However, it still does so imperfectly, with variations in accuracy across regions, suggesting that there are still unexplained factors contributing to the regional production parameters. Fig. 4c shows longitudinal predictions in regions of interest for a single subject with five scans (the greatest number of scans in the cohort). With the exception of the right entorhinal cortex, all data points shown are captured within the confidence intervals, including the average increase across regions. The same predictions are shown for the remaining subjects in Fig. S2 and all show a similar level of accuracy. These results demonstrate the power of a simple model with two free parameters in accurately predicting the regional progression of a individual *τ*P PET progression that may provide benefit for clinical researchers.

## 3 Discussion

We have derived a physics based generative model to describe *τ*P PET data in terms of underlying *τ*P dynamics and applied it data from ADNI and BF2 to understand how it compares to other models present in the literature and how it can inform us about *τ*P transport and production dynamics in the human brain. We have shown that this model can predict accurately longitudinal regional *τ*P PET progression in AD subjects. Furthermore, by performing inference across different patient groups across the AD disease timeline, we uncover temporal changes in transport and production dynamics, showing an initial transport dominated phase associated with primary tauopathy and seeding, followed by an accelerated production dominated phase indicative of secondary tauopathy. There have been several studies proposing different models of proteopathy in AD, a key difference among them being descriptions of the *τ*P production process, which vary widely in complexity [22, 28, 24, 37, 27, 25, 26]. Here we present a parsimonious model of *τ*P transport that relies on regionally specific carrying capacities and we showed through model selection, Section 2.2, that it is able to outperform other models proposed in the literature. A possible cause of regional heterogeneity in carrying capacities is heterogeneity in regional risk factors that promote *τ*P proliferation, the most likely of which is A*β*. A*β* has its own spatial topography within AD patients, being particularly present throughout the fronto-partietal-temporal default mode network and stimulating neuronal hyperactivation [11, 38, 39]. The presence of A*β* will have a two-fold effect on *τ*P dynamics, first through a catalysing effect on the production of *τ*P [40, 41, 42] and second by promoting the activity dependent spread and production through the functional networks [43, 44]. In Thompson et al 2020, we formulated a model describing the dynamic interaction between A*β* and *τ*P, that predicts an increase in carrying capacities based on A*β* concentration [45], however, further work toward simplifying the model will be necessary before it can be used for inference with patient data.

Another key set of factors contributing to regional vulnerability are genetic markers. It has already been shown in mice models of AD that gene expression patterns can inform spreading of *τ*P [31] and human modelling of Parkinson’s disease has shown how gene expression patterns can inform regional vulnerability to create a model of toxic protein spread in Parkinson’s disease [46]. There are several candidate genes for modelling regional vulnerability in AD, most notably microtubule association protein tau (MAPT), as a proxy for relative baseline *τ*P vulnerability [47] and apolipoprotein-E (APOE) for those patients with the APOE*ϵ*4 mutation [29, 48, 49]. While there are many other candidate genes that may influence regional vulnerability, care should be taken to avoid creating overdetermined models. In sum, while the work here provides compelling evidence for the necessity of regional vulnerabilty, further work should seek to explain the mechanisms through which regional carrying capacities emerge from a culmination of regional risk factors, such as A*β* deposition and gene expression patterns.

There is extensive evidence of *τ*P transport and production throughout the brain, however, it has not yet been determined whether one of these processes dominate the other and whether their relative contributions to disease progression changes over time. To this end, we sought to determine whether inferred parameters of our model change in groups across the disease timeline. We find that during early stages of disease, when there is a low *τ*P concentration in the medial temporal lobe, *τ*P dynamics are transport dominated but become production dominated later in disease. This supports previous work by Meisl et al. [27] who show through an analysis of multiple datasets and methods of *τ*P quantification that *τ*P dynamics are production dominated from Braak stage 3 onwards. This is consistent with our work, considering individuals who are positive on tau-PET in early Braak stage regions may already show fairly advanced Braak stages at autopsy [50] analogous to individuals at middle Braak stages used by Meisl et al. 2021. These results suggest that, in early AD, *τ*P seeds invade connected regions from the medial temporal lobe, but the overall concentration does not grow substantially. Only in later stages of AD is the spread production-dominated and driven by fast increases in concentration gradients, leading to progressive Braak-like staging. These results also support the utility of the logistic production model in being able to describe longitudinal A*β*^+^/*τ*P^+^ data (Fig. 2 & Fig. S1), since seeds would already be densely present around the cortex and progression is driven by *τ*P production. The results are also consistent with experimental evidence showing that *τ*P seeds are present before *τ*P pathology [20, 36].

Together the results are indicative of an intrinsically spatiotemporal process, with a primary, transport dominated tauopathy resulting in *τ*P seeds spreading from the medial temporal lobe to axonally connected regions, followed by a secondary, production dominated tauopathy, likely resulting from A*β* interaction, resulting in fast regional accumulation and sequential, Braak-like staging. This is in slight contrast to the largely temporal process of A*β*, described by [51], where A*β* begins as being diffusely present throughout the brain, but increases in concentration at different rates due to regional vulnerabilities.. These results suggest that the early period of AD during which tau is more readily transported between brain regions may be a critical time for intervention. Many immunotherapies currently being developed act on extracellular tau [52] and should therefore interrupt tau transmission through the extracellular space of the synaptic junction. If AD is a consequence of first tau spread and then production, it will be crucial for these immunotherapies to be administered early in the AD process to halt the widespread transmission of tau before accelerated local production can occur. In contrast, therapies that act to reduce intracellular tau concentration should be effective in slowing AD progression across the AD continuum, regardless of whether widespread tau transmission has occurred [53].

This work presents a step forward in whole brain *τ*P modelling, however, there are still many obstacles that are not addressed here. First, there are limitations that pertain to the sparsity of longitudinal data. In this work, we fix a number of parameters to ensure the practical identifiability of the models given the available data. In particular, we fix baseline values and carrying carrying in the dynamical system, and subject initial conditions in the probabilistic model. By fixing baseline values and carrying capacities, we are unable to determine whether these also undergo dynamical changes. This also limits the direct application of the model to other tauopathies that exhibit different tau PET profiles. This will, in part, be addressed by including dynamical regional risk factors, however, the time over which changes in these factors can be observed will remain limited until more longitudinal data is available. Second, there are limitations related to the scale at which we are modelling. While the model we present here is derived from a physics-based model, the model reduction comes at the cost of a loss in mechanistic insight into precise transport and production mechanisms. This will remain a hard limitation while we work with macroscale brain data. In addition, tau PET data is intrinsically limited by resolution, inability to detect early changes, and non-specific and off-target binding sources, collectively providing a source of uncertainty that affects parameter identifiability of intricate processes such as transport. Therefore, while the modelling results suggest changes to transport and production across the AD continuum, our conclusions are limited by the nature of PET measurements and require experimental validation. A potential avenue to address this limitation will be the development of multiscale models that rely on in-vitro or animal studies for calibration and permit macro-scale reduced order models.

The primary contribution of this work has been to supply a parsimonious account of regional *τ*P dynamics in AD. Future work should seek to build upon this, adding more information and data to probe the unexplained dynamics in AD. Most pressingly, these include dynamical interactions between A*β* and *τ*P in a sufficiently simple way to accommodate the ability to perform inference with patient data. Furthermore, the study sheds light on potential avenues of clinical investigation into anti-tau therapies by showing how targeting different tau processes (transport or local production) at different times during the AD continuum may be essential for effective intervention.

## 4 Methods

### 4.1 Data Processing

We use PET from the Alzheimer’s Disease Neuroimaging Initiative (adni.loni.usc.edu). ADNI is a public-private partnership with the aim of using serial biomarkers for measuring the progression of AD. For up-to-date information, see www.adni-info.org. We download the fully processed *τ*P PET and magnetic resonance image (MRI) data, summarised as standardized uptake value ratios (SUVR) and volumes for each of the regions in the Desikan-Killiany-Tourville (DKT) atlas. We re-normalise individual SUVR using an inferior cerebellum SUVR reference region. Amyloid status for ADNI subjects was also downloaded from ADNI and used to classify subjects.

We also use data from BioFINDER2 study, which uses the RO948 *τ*P PET radiotracer. The *τ*P data are analysed using a the analysis pipeline detailed in [54]. Four subjects from the BF2 A*β*^+^/*τ*P^*−*^ group are removed due to high off-target binding in the skull/meninges or MRI registration issues. Amyloid status was determined by Gaussian mixture modelling as detailed in [55, 56]. In both cohorts we select only subjects who have at least three *τ*P PET scans to allow for inference on the time-series model. ADNI and BF2 *τ*P PET data are summarised in table 3.

**Table 3:** Demographics for ADNI and BF2 cohorts.

|  | Group | N. Subjects | Age | Female | Education | CN | MCI | AD |
| --- | --- | --- | --- | --- | --- | --- | --- | --- |
| ADNI | $A\beta^+/\tau P^+$ | 31 | 74.6 | 0.61 | 16.45 | 0.39 | 0.45 | 0.16 |
| | $A\beta^+/\tau P^-$ | 29 | 78.6 | 0.41 | 16.59 | 0.51 | 0.45 | 0.03 |
| | $A\beta^-$ | 59 | 73.1 | 0.54 | 16.71 | 0.63 | 0.36 | 0.01 |
| BF2 | $A\beta^+/\tau P^+$ | 54 | 73.9 | 0.65 | 11.6 | 0.2 | 0.43 | 0.37 |
| | $A\beta^+/\tau P^-$ | 18 | 72.1 | 0.5 | 13.2 | 0.61 | 0.31 | 0.08 |
| | $A\beta^-$ | 53 | 66.9 | 0.5 | 12.8 | 0.81 | 0.19 | 0.0 |

We also use data from the Swedish BioFINDER-2 study (NCT03174938), which uses the RO948 *τ*P PET radiotracer. All participants were recruited at Skåne University Hospital and the Hospital of Ängelholm, Sweden and the cohort covers the full spectrum of AD, ranging from cognitively normal individuals, patients with mild cognitive impairment (MCI) and with dementia. All details about the cohort have been described previously [54]. Amyloid status was determined by amyloidPET (flutemetamol) based on a previously established cutoff from Gaussian mixture modelling as detailed in [55]. Patients with dementia do not undergo amyloid-PET and thus amyloid status was based on the CSF A*β* 42/40 ratio [56]. The *τ*P data are analysed using the analysis pipeline detailed in [54]. Briefly, SUVR images were generated using the inferior cerebellum as reference region and average SUVR were extracted from the DKT atlas. Four subjects from the BF2 A*β*^+^/*τ*P^*−*^ group are removed due to high off-target binding in the skull/meninges or MRI registration issues. In both cohorts we select only subjects who have at least three *τ*P PET scans to allow for inference on the time-series model. ADNI and BF2 *τ*P PET data are summarised in table 3.

We perform inference over three groups: A*β*^*−*^, A*β*^+^/*τ*P^*−*^, A*β*^+^/*τ*P^+^ We distinguish between *τ*P^*−*^ and *τ*P^+^ using a *τ*P PET SUVR cut-off for two composite regions, as detailed in [33]. There are two cut-offs, one for determining *τ*P positivity in the medial temporal lobe (MTL, defined as the mean of the bilateral entorhinal and amygdala), and another for neocortical positivity (defined as the middle temporal and inferior temporal gyri). The thresholds for the composite regions are based on regional Gaussian mixture models, as previously described [24]. For each composite, we average the SUVR values from the constituent regions and fit a two component Gaussian mixture model. The threshold for the region is then set to the SUVR at which there is a 50% chance of being *τ*P^+^. For ADNI, the thresholds are 1.375 and 1.395 and for BF2 they are 1.248 and 1.451 for the MTL and cortical composites, respectively. We define a subject as being *τ*P^+^ if their last scan is suprathreshold in either the MTL or cortical *τ*P PET SUVR and *τ*P^*−*^ if the SUVR value is below both SUVR thresholds.

### 4.2 Mathematical Models of proteopathy

### 4.3 Structural Connectome Modelling

We use the structural connectome to model the transport of *τ*P between brain regions. To generate structural connectomes, we use diffusion weighted MRI images of 150 healthy individuals from the Human Connectome Project (HCP) [57, 58]. From these data, connectomes are derived using the probabilistic tractography algorithm probtrackx [59], available in FSL, using 10000 samples per voxel, randomly sampled from a sphere around the voxel centre. The number of streamlines between each of *R* regions in the DKT atlas are summarised as an adjacency matrix, **A**, that defines our connectome graph, *G*. To model transport of *τ*P between regions, we use the graph Laplacian of *G*, given by:

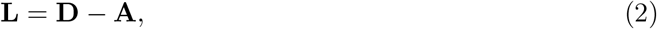

where **D** is the degree matrix, **D** = diag(**A** · **1**). To ensure a transport process respects mass conservation across regions of varying volumes, we weight the graph Laplacian by regional volumes,

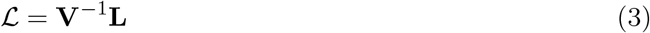

where **V** = diag (**v***/v*_*r*_), and **v** = (*v*_1_, *v*_2_, … *v*_*R*_) is a vector of regional volumes and *v*_*r*_ is a reference region. In Section 2, we model three groups of subjects, A*β*^*−*^, A*β*^+^/*τ*P^*−*^ and A*β*^+^/*τ*P^+^, to reflect changes in volume across the disease timeline and variation in individual brain volumes, we define **v** and *v*_*r*_ on a group and individual level, respectively. For a given group with *N* subjects, we define the normalised volume of a region *v*_*i*_ as

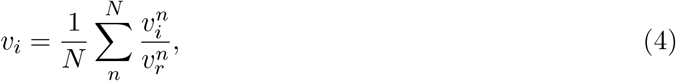

taking 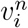 as the initial volume of the *i*th region and *n*th subject and 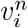 regional volume for the *n*th subject. Then **v** is the average normalised volume per subject in a cohort.

Using the graph Laplacian, we define the *diffusion model*,

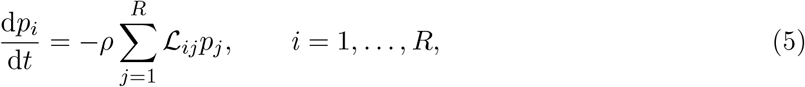

where *p*_*i*_ is the protein concentration at node *i* and *ρ* is a transport coefficient.

### 4.4 Local model of Tau Proliferation

We start with a coupled model of healthy and toxic protein, the heterodimer model, from which we aim to derive a simplified model of toxic protein dynamics that includes regional information. The heterodimer model on a network is:

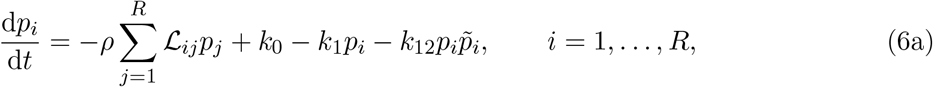

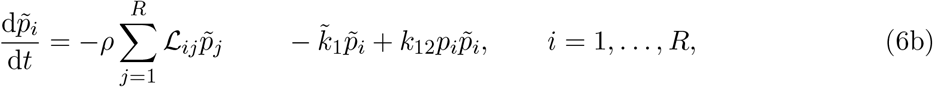

where *p*_*i*_, 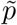 are, respectively, the healthy and toxic protein concentration at node *i, k*_0_ is the natural production rate of healthy protein, *k*_1_ and 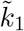 are the clearance rates of healthy and toxic proteins, respectively, and *k*_12_ is the rate of conversion from healthy proteins into toxic proteins [35]. To simplify the heterodimer model, we can follow a similar procedure to that presented in [35], by linearising around a healthy state. Assuming an homogenous state with 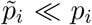, implies 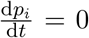 and 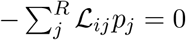 for *i* = 1 … *R*. Then, linearising around 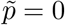, we have

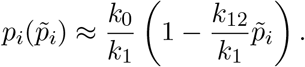

Substituting this expression for *p*_*i*_ into equation (6b) we obtain,

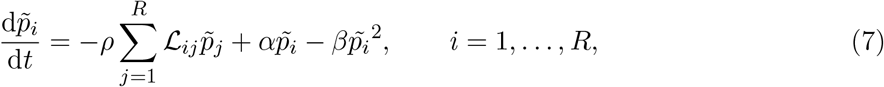

where

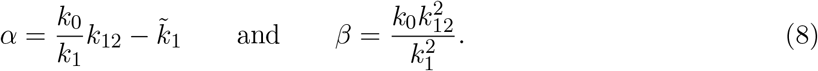

To derive the canonical FKPP model, we perform a change of variables into 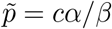, rescaling by the carrying capacity giving:

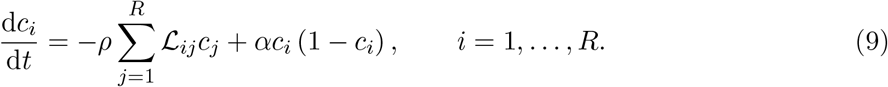

In practice, we introduce a new variable *p*_*∞*_ = *α/β*, to obtain the *global FKPP model:*

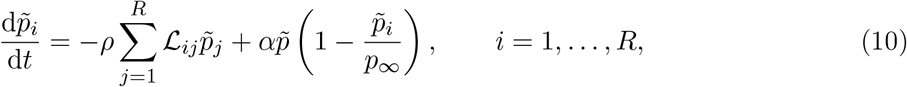

and set *p*_*∞*_ to be the maximum regional carrying capacity inferred from mixture modelling. This ensure the data used for fitting are the same across models.

Next, we derive a model for SUVR concentration *s*_*i*_ with regional carrying capacities and baseline values to model with the requirement that at node *i*, the healthy state corresponds to a baseline value of *s*_*i*_ = *s*_0,*i*_ and the fully toxic state has asymptotic value *s*_*i*_ = *s*_*∞,i*_ as *t* → ±∞. This model is a small generalisation of Eq. (7) that amounts to take *α* and *β* as regionally dependent. It takes the form of the *local FKPP model* :

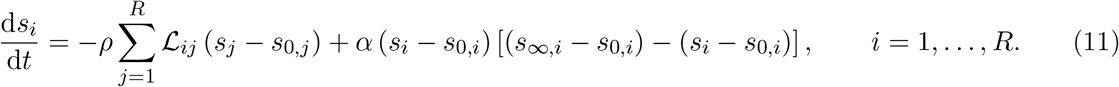

The *logistic model* is then simply obtained by taking *ρ* = 0:

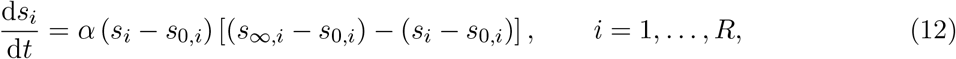

where at each node the variable *s*_*i*_ connects asymptotically, for *t* → ±∞, the healthy state *s*_0,*i*_ to the toxic state *s*_*∞,i*_ (which implies there is no mechanism for propagation from node to node in this model).

### 4.5 Estimating fixed model parameters

To estimate the fixed parameters for **s**_0_ and **s**_*∞*_, we fit a two component Gaussian mixture model to population level data of regional SUVR. For regions in which a reliable measure of *τ*P SUVR can be obtained, we expect to see two separate distributions, a *τ*P^*−*^ distribution capturing the expected *τ*P load in a given region, and a *τ*P^+^ distribution describing the pathological *τ*P load [24]. Using the fitted Gaussian mixture models, we approximate **s**_0_ as the mean of the *τ*P^*−*^ distributions and **s**_*∞*_ as the 99-th percentile of the *τ*P^+^ distributions. For subcortical regions it is not possible to obtain reliable *τ*P PET signal due to off-target binding [60, 61] and we therefore exclude these regions from our model, leaving a total of 72 regions. For ADNI, we use the multi-tau cohort of AV1451 PET data, detailed in [24]. For BF2 data, we use all available RO948 PET scans.

### 4.6 Probabilistic model

For each of the three groups, A*β*^*−*^, A*β*^+^/*τ*P^*−*^, A*β*^+^/*τ*P^+^, we use a hierarchical model, factoring over patients and scans. In each group there are *N* subjects, each of whom have *T*_*n*_ scans, for *n* = 1 … *N* subjects, summarised over *R* regions, (*R* = 72). The observations times, i.e. scan dates, are denoted by 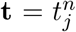 for *j* = 1 … *T*_*n*_, *n* = 1 … *N*. We denote the full data set for a group as **Y** and individual subject data as 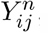, corresponding to the *n*th subject, at scan *j* and region *i*. For a single subject, we have the following data generating function:

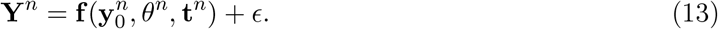

where **Y**^*n*^ is the individual data for *R* regions and *T*_*n*_ time points, with initial conditions **y**_0_, model parameters *θ*, and observations times **t**. The data are generated by a dynamical systems, **f**, with observation error *ϵ*. To derive a likelihood function from Eq. (13), we assume the observations errors are independently and identically distributed and sampled from a Gaussian distribution with standard deviation *σ*. The data generating distribution for a single observation from a subject is then:

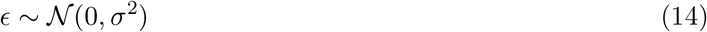

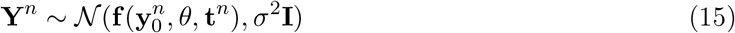

To extend this to a hierarchical population model, we define random variables, 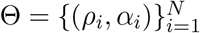, encoding subject specific model parameters and hierarchical population parameters, Ω = {*ρ*_*μ*_, *ρ*_*σ*_, *α*_*μ*_, *α*_*σ*_}, upon which each Θ_*i*_ depends. In Sections 2.2 and 2.3, we assume fixed initial conditions, 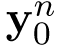, and observations times, **t**^*n*^, taken as the first *τ*P PET scan and scan dates respectively. Then the likelihood function for a single subject under the hierarchical model is:

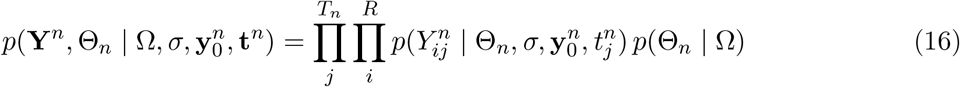

where the first term inside the product on the right hand side is the contribution of the subject level model and the second term is the hierarchical model. Then the posterior for all subjects, hierarchical parameters, subject specific parameters and observation noise is:

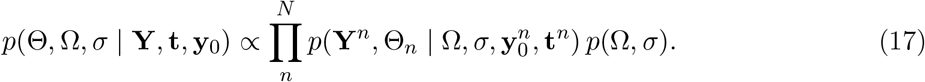

### 4.7 Inference Algorithm

We run inference for each patient group separately using a Hamiltonian Monte Carlo No-U-Turn Sampler (NUTS) to sample from the group posterior distribution. We use the same priors across patient groups, provided in table 4. We use weakly informative priors based on scales at which we expect to observe parameter values and ensure the transport parameter is positive. The NUTS sampler is initialised with a unit diagonal Euclidean metric and a target acceptance ratio of 0.8. For each patient group, we collected four chains each with 2000 samples. All chains showed good convergence (measured by 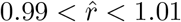) with no post warm-up numerical errors associated with the NUTS sampler.

**Table 4:**
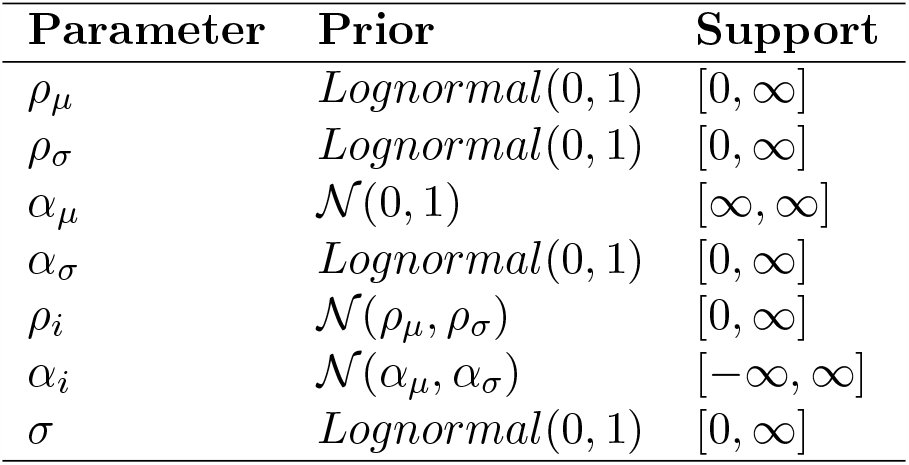
Prior distributions for hierarchical model parameters.

### 4.8 Model Assessment

In Section 2.2 we use two metrics to compare a family of models, the maximum log-likelihood and the expected log predictive density (ELPD). The maximum log-likelihood is used to compare the in-sample accuracy of each of the model’s fit to the data using a NUTS sampler. The maximum log-likelihood estimate is taken as the maximum log-likelihood from the posterior samples, from inference on all available patient data.

The ELPD is used for estimating the out-of-sample predictive accuracy and is adapted from [62]. To do this, we use A*β*^+^/*τ*P^+^ subjects who have more than three *τ*P PET scans (N = 10), using only the first three scans for training and remaining scans to measure predictive accuracy. For our model, the ELPD is then calculated as:

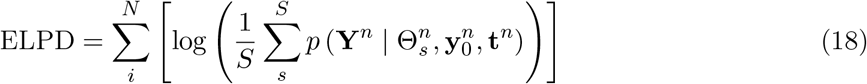

where **Y**^*n*^ are the unobserved data, 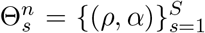 are posterior samples of model parameters, 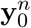 are subjects initial condition and **t**^*n*^ are scan dates, each for *n* = 1 … *N* subjects.

The final method we use for model assessment is the comparison to shuffled data to examine whether the posterior distributions generated inferred from the true data are due to meaningful *τ*P signal or statistical properties of the data. We perform this test on the A*β*^+^/*τ*P^+^ and A*β*^+^/*τ*P^*−*^ groups by random spatial shuffling of the data. The same random permutation are applied to the regional baseline values and carrying capacities. The inference algorithm in Section 4.7 was then applied to the shuffled dataset and 1000 posterior samples were collected. This process was repeated 10 times for each group. In the A*β*^+^/*τ*P^+^ positive group, one chain failed to converge and was discarded.

## Acknowledgement

The work of P. Chaggar was supported by funding from the Engineering and Physical Sciences Research Council grant EP/L016044/1 and F. Hoffmann-La Roche. Author J. Vogel was supported by the SciLifeLab and Wallenberg Data Driven Life Science Program (grant: KAW 2020.0239). A Pichet-Binette is supported by a postdoctoral fellowship from the Fonds de recherche en Santé Québec (298314). S. Jbabdi is supported by Wellcome SRF (221933/Z/20/Z) and Wellcome Collaborative Award (215573/Z/19/Z). G. Klein and S. Magon are supported by F. Hoffmann-La Roche. The BioFINDER study was supported by the Alzheimer’s Association (SG-23-1061717), Swedish Research Council (2022-00775), ERA PerMed (ERAPERMED2021-184), the Knut and Alice Wallenberg foundation (2017-0383), the Strategic Research Area MultiPark (Multidisciplinary Research in Parkinson’s disease) at Lund University, the Swedish Alzheimer Foundation (AF-980907), the Swedish Brain Foundation (FO2021-0293), The Parkinson foundation of Sweden (1412/22), the Cure Alzheimer’s fund, the Konung Gustaf V:s och Drottning Victorias Frimurarestiftelse, the Skåne University Hospital Foundation (2020-O000028), Regionalt Forskningsstöd (2022-1259) and the Swedish federal government under the ALF agreement (2022-Projekt0080). The precursor of 18F-flutemetamol was sponsored by GE Healthcare. The precursor of 18F-RO948 was provided by Roche. The work of A. Goriely was supported by the Engineering and Physical Sciences Research Council grant EP/R020205/1. The funding sources had no role in the design and conduct of the study; in the collection, analysis, interpretation of the data; or in the preparation, review, or approval of the manuscript.

Data collection and sharing for this project was funded by the Alzheimer’s Disease Neuroimaging Initiative (ADNI) (National Institutes of Health Grant U01 AG024904) and DOD ADNI (Department of Defense award number W81XWH-12-2-0012). ADNI is funded by the National Institute on Aging, the National Institute of Biomedical Imaging and Bioengineering, and through generous contributions from the following: AbbVie, Alzheimer’s Association; Alzheimer’s Drug Discovery Foundation; Araclon Biotech; BioClinica, Inc.; Biogen; Bristol-Myers Squibb Company; CereSpir, Inc.; Cogstate; Eisai Inc.; Elan Pharmaceuticals, Inc.; Eli Lilly and Company; EuroImmun; F. Hoffmann-La Roche Ltd and its affiliated company Genentech, Inc.; Fujirebio; GE Healthcare; IXICO Ltd.; Janssen Alzheimer Immunotherapy Research and Development, LLC.; Johnson and Johnson Pharmaceutical Research and Development LLC.; Lumosity; Lundbeck; Merck and Co., Inc.; Meso Scale Diagnostics, LLC.; NeuroRx Research; Neurotrack Technologies; Novartis Pharmaceuticals Corporation; Pfizer Inc.; Piramal Imaging; Servier; Takeda Pharmaceutical Company; and Transition Therapeutics. The Canadian Institutes of Health Research is providing funds to support ADNI clinical sites in Canada. Private sector contributions are facilitated by the Foundation for the National Institutes of Health (www.fnih.org). The grantee organization is the Northern California Institute for Research and Education, and the study is coordinated by the Alzheimer’s Therapeutic Research Institute at the University of Southern California. ADNI data are disseminated by the Laboratory for Neuro Imaging at the University of Southern California.

Data were provided (in part) by the Human Connectome Project, WU-Minn Consortium (Principal Investigators: David Van Essen and Kamil Ugurbil; 1U54MH091657) funded by the 16 NIH Institutes and Centers that support the NIH Blueprint for Neuroscience Research; and by the McDonnell Center for Systems Neuroscience at Washington University.

## Data Availability

Pseudonymized data from the BioFINDER study can be shared by request from a qualified academic as long as data transfer is in agreement with EU legislation on the general data protection regulation and decisions by the Swedish Ethical Review Authority and Region Skåne, which should be regulated in a material transfer agreement.

## Code Availability

Code for the analysis and visualisations produced in this manuscript are available at https://github.com/PavanChaggar/local-fkpp. Analysis was conducted using the Julia programming language [63] and we make particular use of DifferentialEquations.jl [64], Turing.jl [65] and Makie.jl [66].

## 5 Supplementary Information

**Figure S1.**
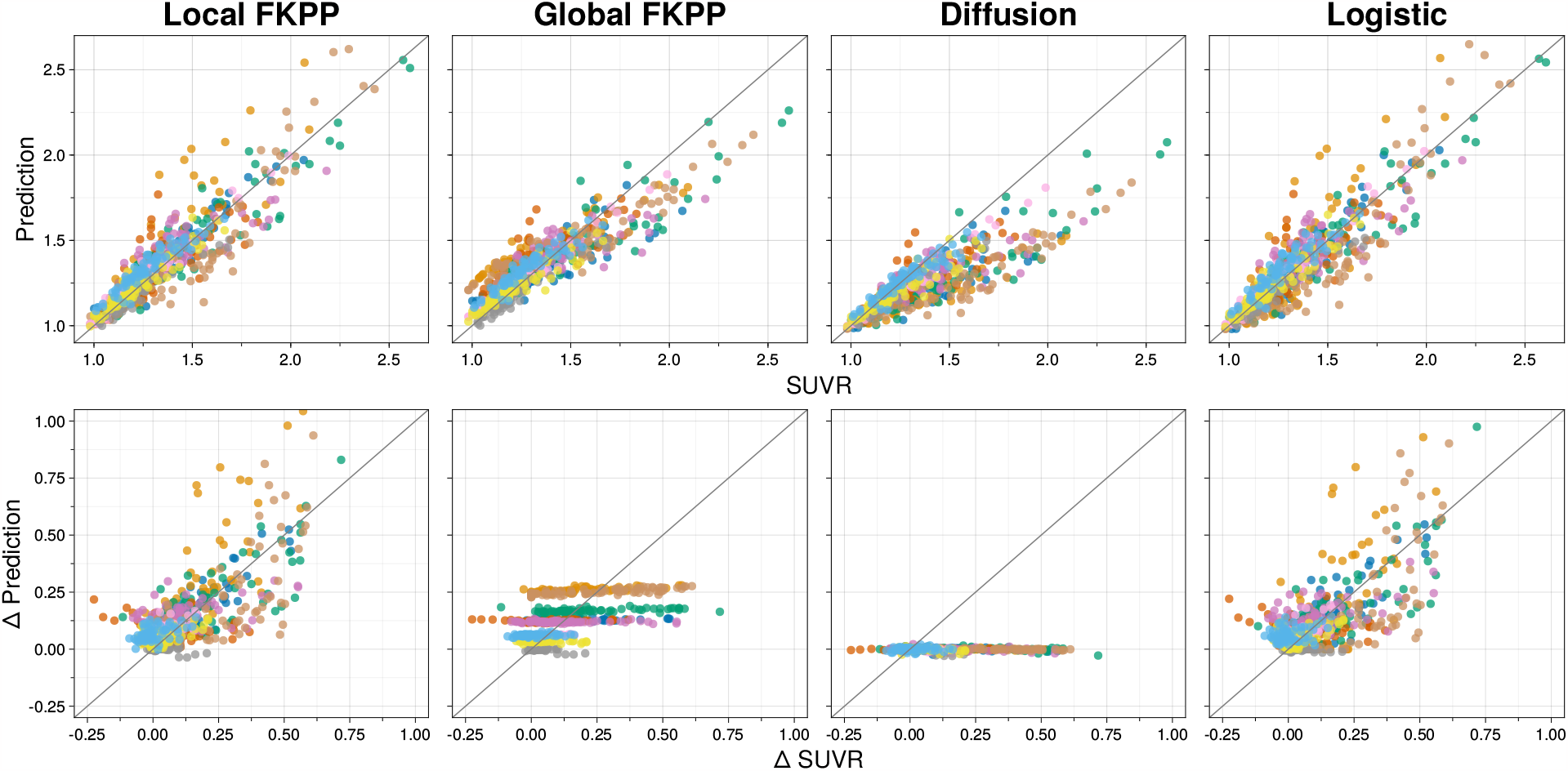
Out-of-sample fits for four models of proteopathy Top row: predicted vs observed out-of-sample SUVR. Bottom row: predicted change vs observed change from first in-sample scan to last out-of-sample scan. Each color represents a different patient and each point of a given color represents a region.

**Figure S2.**
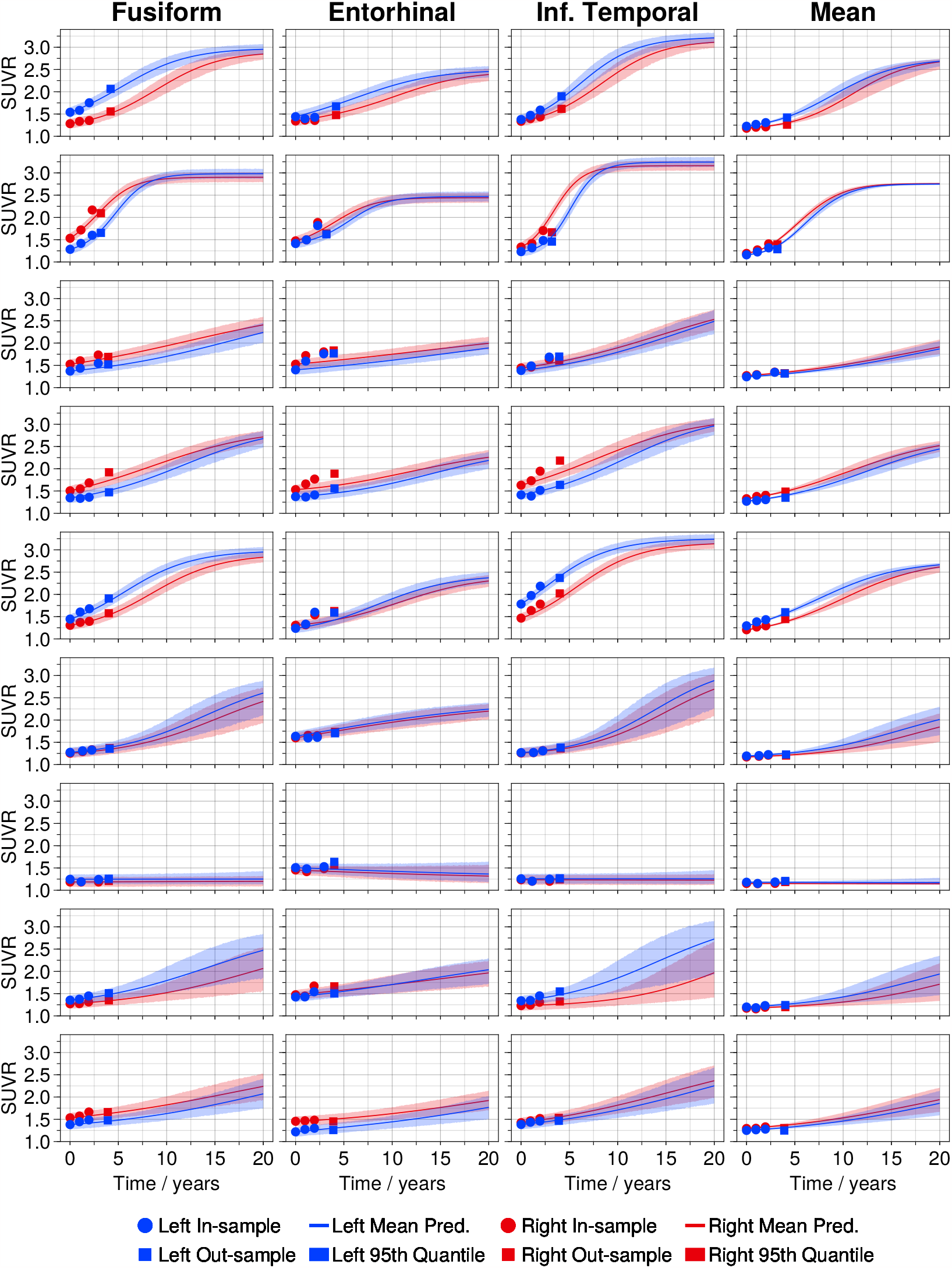
posterior predictive for ADNI A*β*^+^/*τ*P^+^ subjects with at least four scans. Each row is a different subject, each of the first three columns are regions of interest and the final column is the mean SUVR across all regions.

**Figure S3.**
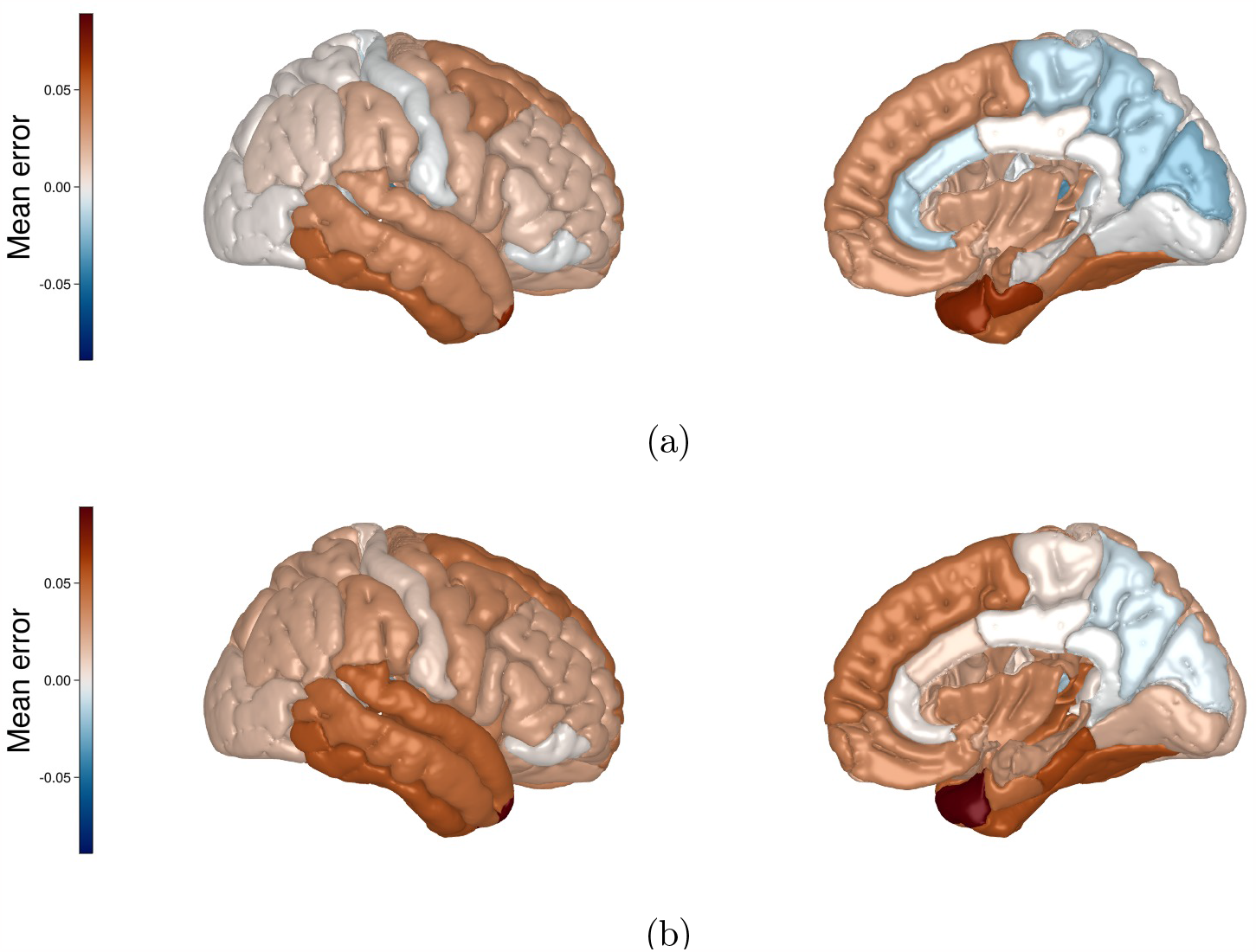
Regional error for in-sample fit against last scan for the A*β*^+^/*τ*P^+^ group. Local FKPP model Fig. S3a and Logistic model Fig. S3b.

